# IMCC: Quantitative Analysis of the Inter-module Connectivity for Bio-network

**DOI:** 10.1101/516237

**Authors:** Pengqian Wang, Jun Liu, Yanan Yu, Yingying Zhang, Bing Li, Dongfeng Li, Wenjuan Xu, Qiong Liu, Zhong Wang

## Abstract

Inter-module connectivity, which tend to connect different communities and maintain network architectural integrity, is contributing to functional coordination and information flow between modules in perturbations. Detecting the strength of inter-module connection is essential to characterize the reactive bio-systematical variation. A quantitative evaluation method for inter-module connections is needed. Here, based on the high-throughput microarray data from mouse, an evaluation approach (named as IMCC) for inter-module connectivity was developed. The IMCC model, which is an integration of direct and indirect inter-module connections, successfully excluded inter-module connections without statistical significance or below the cutoff value, and provided a more comprehensive landscape of inter-module relationships. We showed that the IMCC method reflected a more precise functional coordination between modules according to KEGG database, were validated by topological parameter. Application of IMCC in genome-scale stroke networks deciphered characteristic pathological “core-periphery” structure of modular map and functional coordination module pair.

**Author summery:** Inter-module connectivity, which tend to connect different communities and maintain network architectural integrity, is contributing to functional coordination and information flow between modules in perturbations. Moreover, modular rearrangements provide more efficient ways for phenotype alteration, inter-module connections have been considered to be ‘‘evolutionary interaction switches”. Such modular map rewiring can be used as a network biomarker to characterize the dynamics of drug responses. Detecting the strength of inter-module connection is essential to characterize the reactive bio-systematical variation response to disease or drug. We aim to construct a quantitative evaluation method for inter-module connections. Thus, this study has implications in systematical exploration detailed variation of inter-module pharmacological action mode of drugs.

## 1 Introduction

As accumulated data from high-throughput technologies delineate a holistic view of intra-cellular molecular network, the major challenge in the post-genomic era is deciphering how these entities in the cell work together to execute sophisticated functions [1–4]. The ongoing efforts have been making in decomposing a network into modules [5–9]. This “network dissection” methods may help to reveal modular structure organization in networks. However, inter-module connections, as the ‘backbone’ of cellular networks contributing to functional coordination and information flow between modules in most biological processes, are ever important [10–11]. Such connections tend to connect different communities and maintain network architectural integrity, according to Granovetter’s hypothesis [12–15]. Removal of inter-module connections may lead to collapsed network architecture as well as interrupted information propagation [13]. Moreover, modular rearrangements provide more efficient ways for phenotype alteration [16–17] than genetic variation or modular allostery [18–19], as inter-module edge is more transient and flexible compared with intra-module connections. Inter-module connections have been considered to be ‘‘evolutionary interaction switches” [4, 20–25], because functional innovations often emerge from the rewiring of conserved functional modules to adapt to environment, in the biological system, as one kind of complex adaptive system [26–28]. Such modular map rewiring can be used as a network biomarker to characterize the dynamics of drug responses [29], by identifying and evaluating the drug-conditional existence of collaborations between modules, as a result to reflect the dynamical fluctuations of molecular correlations [30–34]. Therefore, it is an extremely interesting and promising perspective to apply inter-module connectivity analysis to characterize the reactive bio-systematical variation to perturbation, especially the pharmacological mechanism of multiple-target compounds [35–39].

Recently, increasing attention has been paid to inter-module research. Several concepts, such as “bottleneck” [41], “connector” [42], “bridgeness” [42–44], and “fuzzy community” [44–48], have been proposed to investigate the node or module bridging one community to another, regulating the information flow between modules, maintaining dynamic rewiring in networks [41, 49]. The principle of inter-module correlations has been explored and suggested to constitute a form of functional coordination between modules in a pairwise fashion [10, 50]. A set of studies have also focused on the quantitative evaluation method for inter-module connections, such as counting the number of interactions or overlapping nodes [51] between modules or communities, which were commonly conducted to represent as the connections [17, 52–55]. Meanwhile, the significance of inter-module connections has also been calculated by performing hypergeometric test, Fisher’s test [10, 56], Wilcoxon rank-sum test [50], or optimization of the global score [56]. Studies on module coordination have shown that module cooperation should be established not only by direct interactions among modules but also through shared partners [57] or between-modules (or pathways) paths that consist of multiple proteins and interactions [58]. Although previous algorithms proved valuable, it was often ignored that the modules in biological network were of functions, and all of the evaluation methods for inter-module correlations should be oblige to biological functions. Therefore, it is pertinent to introduce biological function to measure reliability and validity of inter-module evaluations. Additionally, inter-module connectivity should be established not only considering direct inter-module connections bridged by shared nodes or interaction, but also through shared partners [57] or between-modular(pathway) path consisting of multiple protein and interactions [58]. To shed light on the inherent connections between modules and consequent inter-module coordination related pathological process and pharmacological mechanism, it is pressed for developing an integrated and more precise evaluation method for quantitative analysis on inter-module connectivity.

In this paper, using genetic interactome and modules based on microarray of MCAO (middle cerebral artery obstruction) mice and WGCNA (weighted gene co-expression network analysis), we proposed a novel approach to evaluate inter-module connectivity, integrating quantitative analysis and statistical significance, named as inter-module connectivity coefficient (IMCC). Also, the biological similarity between modules from the KEGG database were employed to optimized the IMCC model. Then, the IMCC were compared with extant method on real network, based on the precision to reflect the inter-module coordination in KEGG database. And IMCC was also validated by topological parameter. Finally, we applied IMCC in genome-scale stroke networks to decipher characteristic inter-module connectivity related to pathological process.

## 2 Materials and methods

Inter-module connectivity contributed to information propagation and functional coordination between modules. According to local conformations, connections between two modules composed of edges between nodes from distinct modules (direct-edge) or shared nodes from different modules (direct-node) were defined as direct inter-module connections (DIMC), and interactions mediated by genes that associate with both of the two modules were classified as indirect inter-module connections (IIMC).

In order to develop a more comprehensive, objective, and accurate method to measure the module-to-module relationships and reflect the biological coordination between modules, we firstly calculated two types of correlation parameters: SW for DIMC; CT and PS for IIMC; and then we screened these parameters using hypergeometric distribution or cutoff value and integrated the identified parameters; finally, we optimized the integration weight according to KEGG database (Figure. 1).

**Figure 1.**
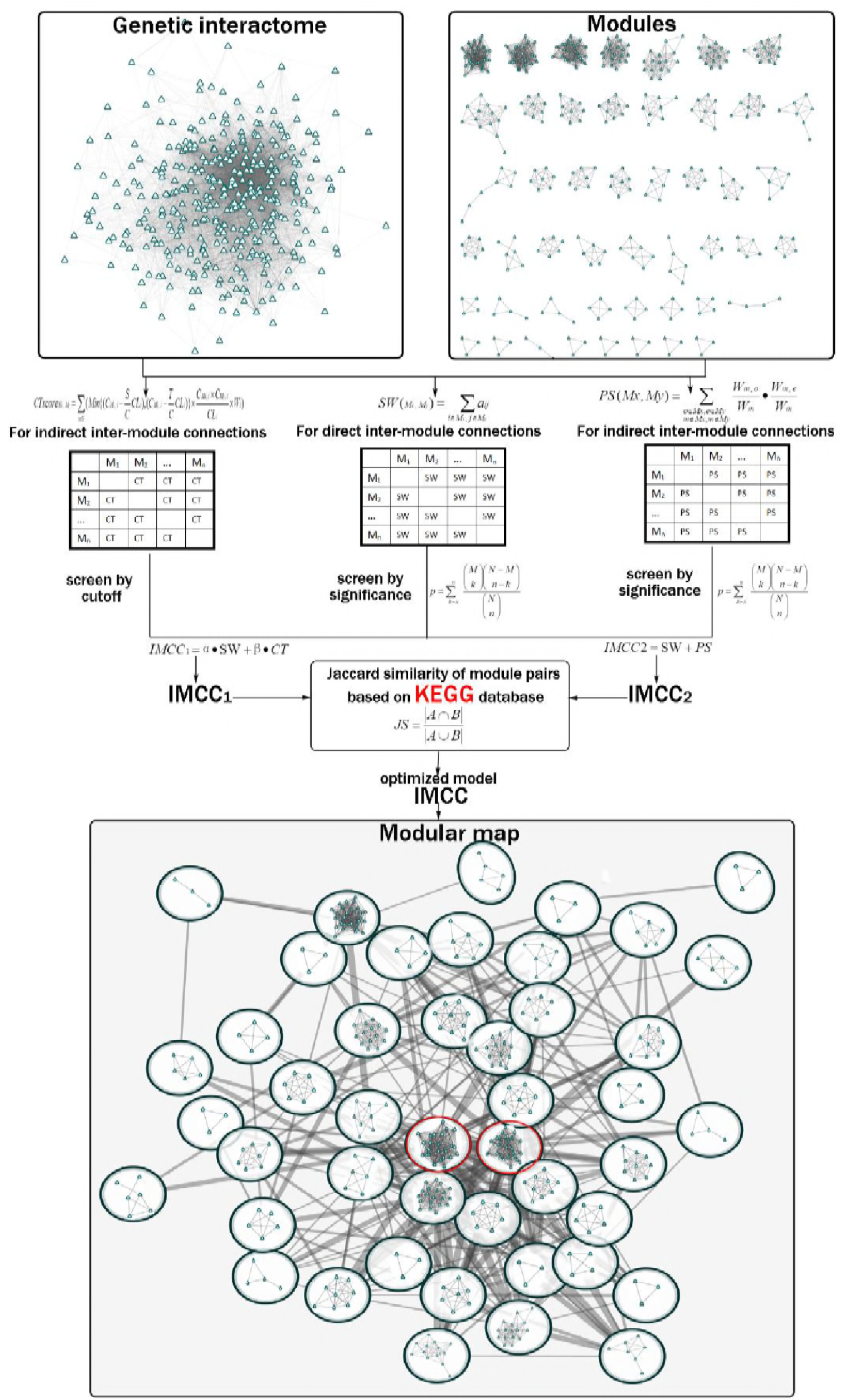
The procedure of computing IMCC model. Based on the genetic interactome and modules originated from microarray of MCAO mice, the parameters for direct or indirect inter-module connections were calculated, screened and integrated. Then the IMCC model were optimized according to KEGG database. The box at the bottom of the diagram was the modular map based on IMCC, in which vertexes denoted modules and the thickness of edges linking pairs of modules was directly proportional to the corresponding IMCC. In this modular map, the red circled modules represented the “core” module pair (module-blue and module-brown).

### 2.1 Parameters calculations

For SW, we calculated the sum of weight of edges between pairs of modules:

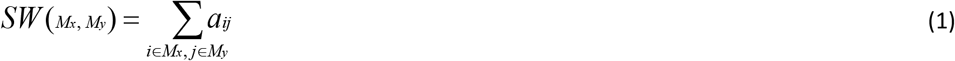

where *M_x_* and *M_y_* denote any two modules connected by at least one edge, *i* and *j* are a gene in *M_x_* and *M_y_*, respectively, and *a_ij_* is the weight of edge between gene *i* and *j*. Using this formula, we calculated the direct inter-module connections for any module pair possessing one or more edges.

To decide whether the inter-module direct connections were statistically significant, we used the P-value of the hypergeometric distribution [11], defined as

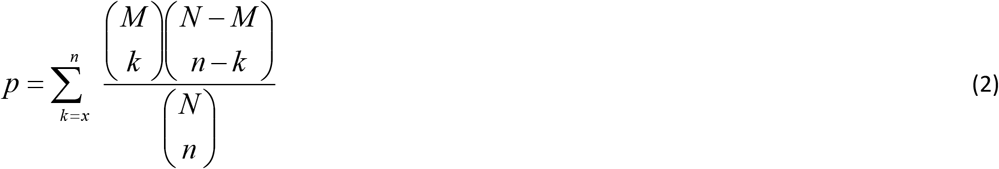

where x is observed inter-module connections, k and n are the numbers of inter-module connections and all possible edges between two modules, respectively; M and N represent the total numbers of inter-module connections and all combinational gene pairs between any two modules in a module to module network, respectively. In this paper, we set 0.05 as P-value threshold. Therefore, inter-module connections are supposed to be present, if P-value ≤ 0.05; and the SW is defined as a valid direct measurement.

For indirect inter-module connections, we introduced two parameters: path strength [58–59] and consistency score [57].

In the light of the network, paths consisted of multiple vertexes and links between them [58]. To simplify the problem, we restricted the length of paths and only considered paths that consist of three nodes (outset (o), mediation (m) and end (e)) with two links.

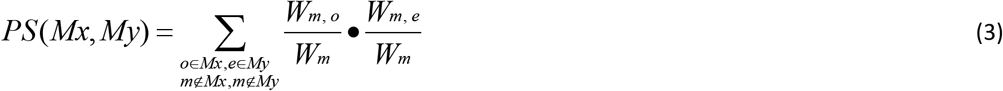

The path strength (*PS*) of a path is defined as the product of the weighted probabilities that mediation chooses outset and end. The weighted probability from *m* to *o* is the ratio of the weight between m and o *(W_m,o_)* to the sum of the weights between m (*W_m_*)and its first neighbors, the same as *m* to *e*.

Hypergeometric distribution was also used to screen the statistically significant PS. However, different from SW, in formula (2), x is observed nodes connecting a pair of module, k and n are the numbers of nodes connecting a pair of modules and all possible nodes connecting the two modules, respectively; M and N represent the total numbers of nodes connecting any pair of module and all possible nodes between any two modules in a module to module network, respectively. In this paper, we set 0.05 as P-value threshold.

We also employed consistency score to measure the inter-module connectivity as described in [57]

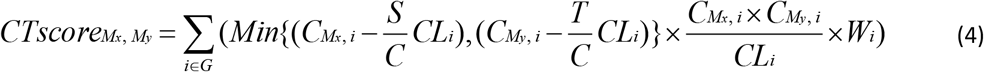

G is a gene set that consists of all genes in network, and C is the total number of genes in G. CLi is the total number of links to gene i; Wi is the weight of gene i in network. S and T are the numbers of genes in modules Mx and My, respectively; CMx,i and CMy,i are the observed numbers of links connecting gene i and modules Mx and My, respectively. Formula (4) was used to compare the weights of genes correlated with a pair of modules with the weights of genes related to only one of the modules [57]. As the CT is a value after comparison with theoretical value, we set cutoff value (10) to screen out the valid CT.

We listed the parameters and screening procedures in Supplementary Table 1 and 2.

### 2.2 Measurement integration

To obtain a more accurate and objective relationship between modules, we merged DIMC and IIMC. Firstly, we set SW of the inter-module connections, whose P-value of hypergeometric distribution was equal to or less than 0.05, as weight of DIMC. In the process of IMCCi integration, SW and CT were two parameters of different dimensions, so it was adopted to correct the two parameters to the value of 0-1. We normalized the two measurements to be numbers between [0–1] by the follow formula:

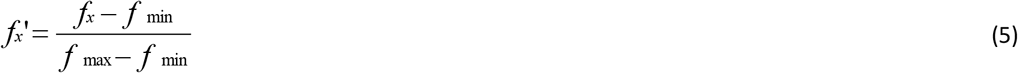

The two parameters were weighted using weighted coefficient α and β for SW and CT, respectively. And the two weighted parameters were summed up, as followed:

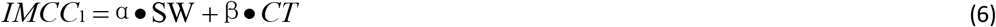

We set α + β=1, and coefficient ratio ρ=α/β. By adjusting ρ value, the effect of SW and CT on IMCC_1_ would be altered. We calculated the IMCC1 when ρ = 1/10, 1/8, 1/4, 1/2, 1/1, 2/1, 4/1, 8/1, and 10/1, respectively.

In the integration of SW and PS, both of them belong to the same dimension, so we plused the two parameters without weighting, defined as:

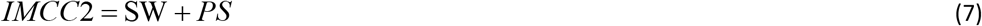

The number of screened SW, CT, and PS were shown in figure 2A and 2B.

**Figure 2.**
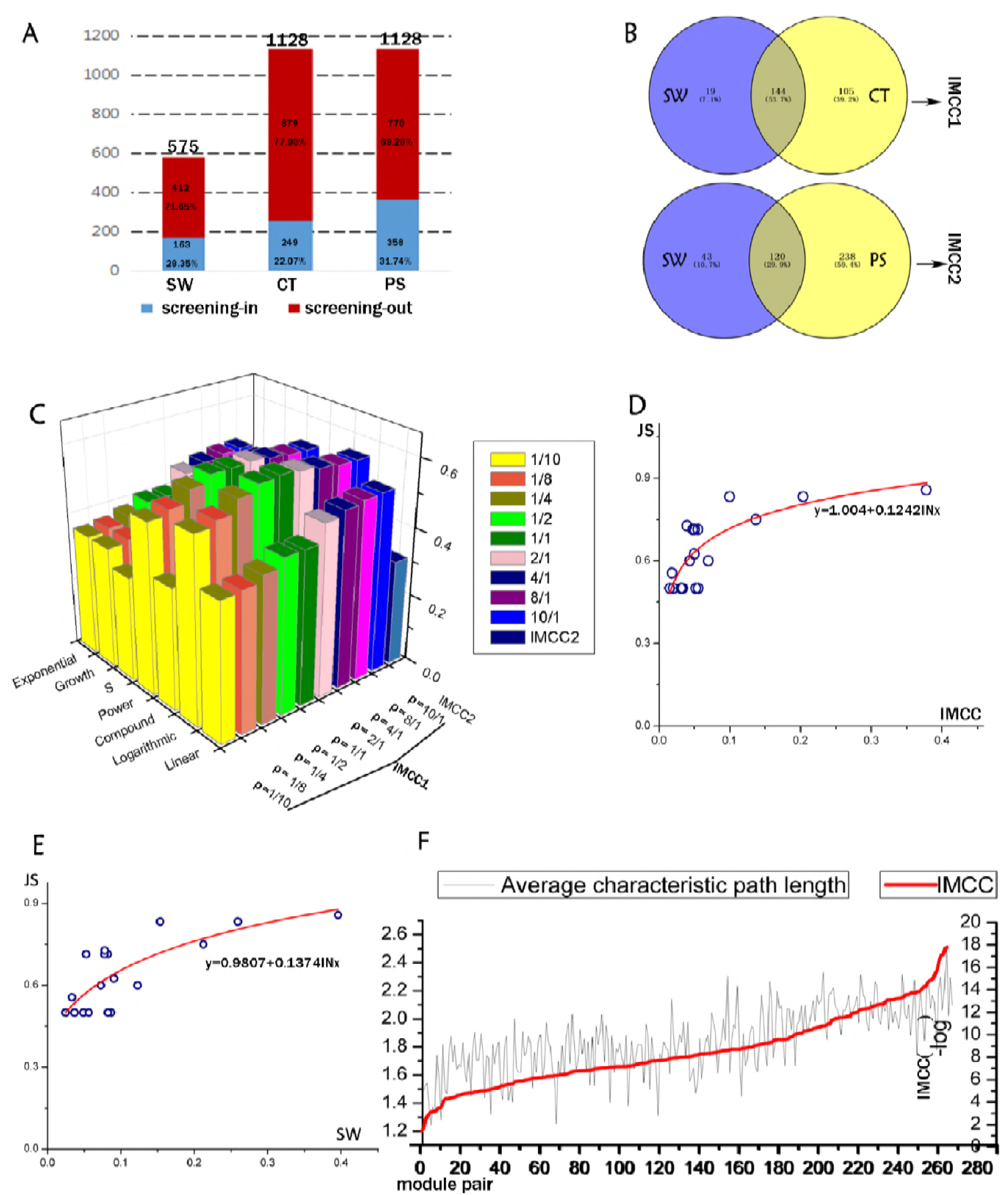
Quantitative evaluation of the IMCC method. A The screening outcomes of the 3 inter-module connectivity parameters (SW, CT and PS). The height of the column represents the total amount of SW, CT and PS, respectively. Red and blue parts of the columns represent the screened out and the remaining parameters, respectively. B The overlapping condition of two parameters to be integrated as IMCC1 and IMCC2. C The R2 of 7 fitting models for IMCC1 with different ρ and IMCC2. D The fitting curves of logarithmic model of JS and IMCC. E The fitting curves of logarithmic model of JS and SW. F The IMCC value against the average characteristic path length of module pairs.

### 2.3 Data sources

We used genetic interactomes and modules constructed based on the microarray datasets of MCAO mice and WGCNA, to illustrate the performance of IMCC method on the inter-module connection calculation.

### 2.4 Data analysis and weighting coefficient optimization

Considering the different characteristics in various types of networks, whether the emphasis of IMCC should be placed on DIMC or IIMC must be consistent with practical applications. In biological networks, the communication between certain modules is commonly mediated by component with important functions; for example, a gene might be a target regulated by two modules competitively. All the inter-module correlations summarized or predicted are presumed to contribute to biological functions. Therefore, it is imperative to introduce biological data to define the best weighting coefficient, in order to select the optimal IMCC. As a result, we employed the KEGG (Kyoto Encyclopedia of Genes and Genomes) database, according to which we calculated the jaccard coefficient of enriched pathways of each module pair.

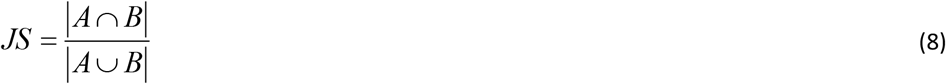

A and B is the category of enriched KEGG terms in module Mx and My, respectively. Therefore, A∩B represents the number of identical categories of KEGG terms between A and B; and A∪B represents the number of all the categories of KEGG terms in both A and B. For example, if A∩B=6, and A∪B=11, then JS will be 0.54545. We presumed that modules enriched with the same KEGG categories might form more dense connections than those with different KEGG categories. We benchmarked the IMCC1 of different ρ values based on JS (Figure 2D, Supplementary Table 3), and the IMCC1 scores were plotted versus the observed JS for each module pair. To obtain more precise results, we removed the outliers (Supplementary Table 3). Through linear and 6 nonlinear curve estimating, we compared the coefficient of determination (R2), and quantitatively identified the best ρ value and the most fitting model (Figure 2C). Our results suggested that the optimal ρ value was 1/1 and the most fitting model was logarithmic model with a R^2^ of 0.616. Thus the final formula for IMCC could be simplified as

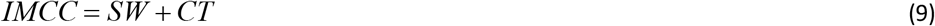

We also plotted the IMCC_2_ against JS, and compared R^2^ of fitted curves of IMCC_1_ and IMCC_2_ to select the optimal integrative method.

### 2.5 Comparison and verification of inter-module average shortest path

For previous algorithms for inter-module were mainly based on the sum of weight of interactions (SW) between pairs of modules, we compared IMCC with SW by plotting the score versus the KEGG accuracy (JS calculated as formula 8), as shown in Figure 2D, E, Supplementary Table 3.

We also compared IMCC with inter-module average shortest path (IMASP), a topological parameter proposed to evaluate the distance of a module pair. IMCC was plotted versus the IMASP, and determination coefficient of curve fitting was also calculated (Figure 2F, Supplementary Table 4).

## 3 Results

In this paper, based on high-throughput microarray data from mouse, we integrated two types of inter-module connections, i.e. DIMC and IIMC, combined with statistical methods, and developed a novel quantitative algorithm termed as IMCC. Using IMCC, we drew more coarse grains from molecular network, and integrated the microscopic molecules into mesoscopic modules.

### 3.1 The IMCC model screens out noise

Biological networks, which were constructed based on a DNA microarray data set and a mathematical model [60], were considered to have high noise due to the false-positive levels inherent in the data set. Taking our experimental network as an example, there were 374 nodes in the co-expression network, whereas most (55.93%) of the shortest path length between any two nodes is gathering in 2, and the third most (13.41%) in 1 (Supplementary Table 5). This indicated that more than 60% of the node pairs could be connected directly or indirectly. Accordingly, in the modular map, the module-module connections also manifested a false-positive property. As for modules, all of the module pairs could be connected directly or indirectly to form a densely interacted modular map. In our data set, the number of direct inter-module connections (DIMCs) was 575, indicating that averagely each module in this map had 24 neighborhood. And the number of indirect inter-module connections (IIMCs) was up to 1128 in the modular map, indicating that any pair of modules was connected by IIMC. In face of such a large number of inter-module connections, how to remove noise interference to accurately screen out the real data about module connections?

Therefore, it seems imperative to screen out the random fluctuations of noise in inter-module connections. We introduced the hypergeometric distribution test (details are described in Methods), which calculates the probability that the specified target is selected from the whole population, so as to identify the valid value of SW (a parameter of DIMC) and PS (a parameter of IIMC) with significance. As the CT (another parameter of IIMC) value is drawn from the comparison with expectation of the whole network, we set the cutoff value of CT at 10 (an inter-module connection with a CT > 10 was considered valid). Comparison of the number of module-module interactions before and after screening revealed large differences. Among the 575 SW, we screened out 412 (71.65%) invalid SW and 163 (28.35%) valid SW remained, according to the P value of hypergeometric distribution. Meanwhile, out of the 1128 indirect module-module interactions, 249 (22.07%) valid CT and 358 (31.74%) valid PS remained after screening based on the cutoff value and hypergeometric distribution, respectively (Figure 2A). After the screening process, we successfully excluded a large number of inter-module connections without statistical significance or below the cutoff value.

### 3.2 The IMCC method provides a comprehensive landscape

The integration of parameters about these direct and indirect connections provides a more complete landscape of inter-module relationships. Two functional modules may bind to each other or target an identical molecule to constitute a competitive regulation, both of which are universally present in biological networks. There were 144 overlapping inter-module connections between SW and CT, and 19 (11.66% of the total SW) and 105 (42.17% of the total CT) specific (non-overlapping) inter-module connections in SW and CT, respectively. Between SW and PS, 120 overlapping inter-module connections were identified, and 43 (26.38% of the total SW) and 238 (66.48% of the total PS) specific inter-module connections were found in SW and PS, respectively (Figure 2B). These findings revealed that a certain number of module pairs were merely correlated by either direct or indirect connections. As a result, our integration model has a wider-range of coverage than solo-direct and solo-indirect inter-module connections, which may provide a more comprehensive landscape of inter-module connections. Details of the integration process for SW and CT, as well as SW and PS were described in Supplementary Table 1 and 2.

### 3.3 The IMCC method reflected inter-module functional coordination

When integrating these parameters, we compared and analyzed two integration models of IMCC (IMCC1 and IMCC2). All the inter-module correlations summarized or predicted are presumed to contribute to biological functions. We employed the KEGG to select the most fitting integration model. By plotting Jaccard Similarity (JS) coefficient of each module pair based on KEGG versus IMCC value, we calculated the precision of IMCC (Supplementary Table 3). Our results (Figure 2C and D) suggested that the optimal ρ value in IMCC1 model was 1/1 and the most fitting model was logarithmic model with a R^2^ of 0.616. In logarithmic model, the R^2^ of different p values showed an obvious peak phenomenon: when ρ=1/1, the R^2^ reached the peak with two sides sloping down to lower values; when ρ=1/10 or ρ=10/1, the minimum R^2^ of each side were observed, respectively (Supplementary Figure 1). Therefore, it is proper to decide that the integrated parameter IMCC1 is more consistent with the KEGG classification than any single index (SW or CT), which would provide more accurate evaluation of the relationship between modules. Using the same method, we plotted the IMCC2 value against JS in KEGG, and found that there was no correlation between the two parameters and the fitting R^2^ was much lower than IMCC1 when ρ=1/1 (Figure 2C, Supplementary Figure 2). Therefore, we chose the IMCC1 model as the final model with a ρ value of 1/1. The final formula can be simplified as follows:

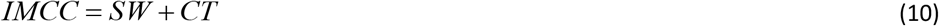

To some extent, the integrated method, which is based on the topology structure of networks, reflects the functional coordination of module pairs.

### 3.4 The IMCC method is more precise and validated by a topological parameter

In the comparison, IMCC showed increased performance of an integrated score relative to the SW. The fitting model of SW was y = 0.1374ln(x) + 0.9807, with R2 equal to 0.580, which is lower than 0.616 of IMCC (Figure 2E). Overall, the results indicated that IMCC achieved better performance on the weighted gene co-expression data. It also means that the results based on IMCC were more consistent with biological function than SW. After all, the IMCC provided a more precise evaluation for inter-module connections.

Unlike protein-protein interaction networks, gene co-expression networks are weighted networks. Thus, inter-module connections in such networks should not only include dichotomous edge (0 or 1), but also quantitative precision information. We introduced the SW, CT and PS to quantitatively analyze these inter-module connections, taking the weight and amount of the edge of molecular networks into consideration. We compared the correlation based on IMCC and a dichotomous topological parameter, inter-module average shortest path (also known as average characteristic path length). The results of the two assessments were generally consistent (Figure 2F), indicating that the two parameters had positive linear correlations with a R^2^ of 0.495 (Supplementary Table 4, Supplementary Figure 3). Therefore, this result successfully validated the IMCC method.

### 3.5 Application of IMCC in genome-scale stroke networks deciphered pathological inter-module connectivity

We applied IMCC on genetic interactomes and modules based on microarray of MCAO mice to explore the characteristic inter-module connections related to stroke. By calculating IMCC between each pair of module, we constructed modular map, in which vertex and edge represent modules and IMCC (as shown in Figure 1). This may contribute in the following two aspects. First, we showed the architecture of the modular map in stroke condition. Second, we analyzed the enriched KEGG pathways of highly correlated gene modules to explore how the pathological pathway interactions contributed to stroke process.

As an overarching view, the modular map exhibited a characteristic “core-periphery” structure (as shown in Figure 1), where the periphery consisted of small well-defined communities, and the core comprised highly interconnected larger modules, which are harder to detect [61–64]. It is suggested that when a biological network is on stress, the inter-module connectivity would decrease [65], to promote this core-periphery structure [63], center of which is conserved and stable module, with crucial role for cell survival rather than development [64, 66], to adapt to the novel situation. In the stroke modular map, the most closely connected modules were a pair of modules (module-blue and module-brown), which can be regarded as “core” presenting characteristic pathological process for cell survival in stroke.

Thus, it is essential to enrich the KEGG pathways of this “core” to explore predominant pathological pathways in stroke. The module-blue was enriched with 12 KEGG pathways; the module-brown was enriched with 4 KEGG pathways (p < 0.05, shown in Table 1). Among these pathways, HTLV-I infection, which were suggested to associate with pathogenic mechanism of immune dysregulation in neuroinflammatory [67], was both enriched in module-blue and module-brown: 5 genes from module-blue and 5 genes from module-brown encoded proteins in this pathway (Figure 3). This indicated that the coordination between tightly connected module pair might be attributed to constituting an identical pathway of biological essence. Furthermore, pathways enriched by two modules showed cross-talk based on the KEGG background. For example, MAPK signaling pathway of module-brown participated in pathways in cancer, long-term potentiation, neurotrophin signaling pathway and foxO signaling pathway of module-blue. Therefore, pathway crosstalk might be responsible for the module coordination. Taken together, such topological inter-module connections turned out to constitute a form of functional coordination between modules. We speculated that this coordination is occurring typically from constituting an identical pathway or forming pathway crosstalk. This may also show the ability of our analysis to detect inter-module connectivity of biological function, which is consistent to prior knowledge and may strengthen the role of module coordination.

**Figure 3.**
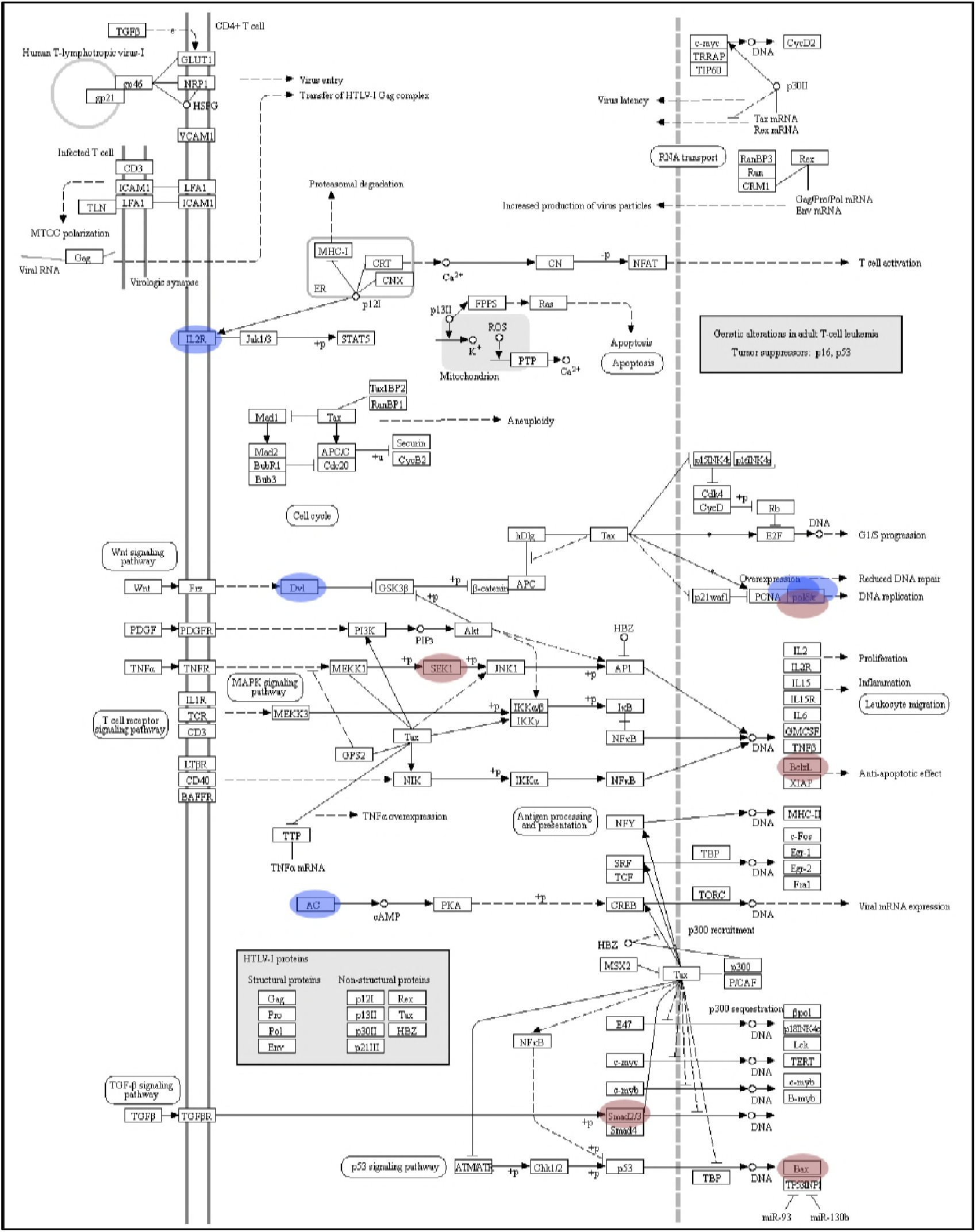
Proteins encoded by genes from module-blue (labelled by blue) and module-brown (labelled by brown) in HTLV-1 infection based on KEGG database.

**Table 1.** Enriched KEGG pathways of module-blue and module-brown.

| Module | Term | P-Value | Genes |
| --- | --- | --- | --- |
| **Module-blue** | Pathways in cancer | 0.001721 | VEGFB, DVL3, TRAF2, ADCY1, BRAF, GNA11 |
|  | HTLV-I infection | 0.003376 | POLD3, DVL3, ADCY1, POLD1, IL2RG |
|  | Long-term potentiation | 0.010084 | ADCY1, BRAF, CAMK4 |
|  | Aldosterone synthesis and secretion | 0.016716 | ADCY1, CAMK4, GNA11 |
|  | Insulin secretion | 0.016716 | ADCY1, GCK, GNA11 |
|  | Cholinergic synapse | 0.027889 | ADCY1, CAMK4, GNA11 |
|  | Neurotrophin signaling pathway | 0.032131 | PDPK1, BRAF, CAMK4 |
|  | Vascular smooth muscle contraction | 0.034593 | ADCY1, BRAF, GNA11 |
|  | FoxO signaling pathway | 0.03816 | PDPK1, BRAF, GADD45A |
|  | Hepatitis C | 0.039204 | TRAF2, PDPK1, BRAF |
|  | Insulin signaling pathway | 0.041326 | PDPK1, BRAF, GCK |
|  | Hippo signaling pathway | 0.047383 | DVL3, BBC3, BTRC |
| **Module-brown** | Amyotrophic lateral sclerosis (ALS) | 1.73E-04 | BAX, GRIN2A, BCL2L1, SOD1 |
|  | MAPK signaling pathway | 0.001913 | RPS6KA4, JUND, MAP2K4, GNA12, HSPA1A |
|  | HTLV-I infection | 0.002702 | BAX, POLD2, MAP2K4, SMAD3, BCL2L1 |
|  | Hepatitis B | 0.040124 | PTK2B, BAX, MAP2K4 |

## 4 Discussion

In this paper, we described an integrative, quantitative approach (IMCC) to quantify inter-module connections. There were several unique features in our methodology: i) Using the statistical screen, the IMCC method presented a novel and powerful tool to filter out the random fluctuations of noise from significant inter-module connections. ii) By integrating the analysis of direct and indirect inter-module connections, we provide a more comprehensive approach for quantification. iii) The biological functions were introduced to evaluated the reliability of inter-module connections. By fitting the IMCC score to the JS of the KEGG category, IMCC is considered a more precise tool for inter-module topology structure analysis reflecting the functional coordination of module pairs, in the comparison with extant parameter. Taken together, this novel model showed good performance to reveal functional interactions among modules.

This capability can be extremely useful for bio-network. In the application in genome-scale stroke networks, we deciphered characteristic pathological “core-periphery” structure and functional coordination module pair. Thus, this method can be attempted to apply in case-control or drug perturbation network to explore detailed variation of inter-module connectivity related to pathological process or pharmacological action. This inter-module connectivity of disease-or drug-conditional existence can be served as network biomarkers to characterize the dynamic response to perturbation. For its flexibility and impact on phenotype alteration, inter-module connection can also be served as targets to design specific drugs. This concept may bring about reinterpreting the connotation of pharmacology. After all, various desirable properties of this algorithm discussed in this work will facilitate the inter-module analysis to be applicable to biological complex systems for disease and drug discovery.

## Supporting information

S1 Text. Supplementary Figures and tables for model and validation. (DOC)

## Acknowledgments

This work was supported by National Science and Technology Major Projects “Major New Drug Development Project”. [grant number 2017ZX09301-059].

The authors declare that they have no competing interests.

